# Masked Minimizers: Unifying sequence sketching methods

**DOI:** 10.1101/2022.10.18.512430

**Authors:** Minh Hoang, Guillaume Marçais, Carl Kingsford

## Abstract

Minimizers and syncmers are sequence sketching methods that extract representative substrings from a long sequence. We show that both these sampling rules are different instantiations of a new unifying concept we call masked minimizers, which applies a sub-sampling binary mask on a minimizer sketch. This unification leads to the first formal procedure to meaningfully compare minimizers, syncmers and other comparable masked minimizers. We further demonstrate that existing sequence sketching metrics, such as density (which measures the sketch sparseness) and conservation (which measures the likelihood of the sketch being preserved under random mutations), should not be independently measured when evaluating masked minimizers. We propose a new metric that reflects the trade-off between these quantities called the generalized sketch score, or GSS. Finally, we introduce a sequence-specific and gradient-based learning objective that efficiently optimizes masked minimizer schemes with respect to the proposed GSS metric. We show that our method finds sketches with better overall density and conservation compared to existing expected and sequence-specific approaches, enabling more efficient and robust genomic analyses in the many settings where minimizers and syncmers are used.

## 1 Introduction

Minimizers [17, 18] are deterministic methods to sample substrings from a sequence at approximately regular intervals such that sufficient information about the identity of the sequence is preserved. Sequence sketching with minimizers is widely used to reduce processing time and memory consumption in bioinformatics programs such as read mappers [9, 12], *k*-mer counters [3, 6] and genome assemblers [20]. Given a choice of *k*-mer length *k* and window length *w* (i.e., a (*w, k*)-window is a substring with exactly *w* overlapping *k*-mers), a minimizer selects the lowest priority *k*-mer from every window in the target sequence according to some total ordering over all *k*-mers. We refer to the set of all *k*-mers sampled in this manner as a minimizer sketch. In many applications, the desirability of a sketch is typically measured by its *density* [14], or formally the ratio between the sketch size and the length of its target sequence.

A low-density minimizer scheme achieves three desiderata: (1) the sketch size is small and will offer significant cost saving to downstream applications; (2) every (*w, k*)-window in the target sequence is represented by at least one *k*-mer in the sketch; and (3) identical windows are represented by the same *k*-mer in the sketch due to the deterministic sampling protocol. It was recently argued that property (2) is not sufficiently robust in sequence alignment applications where mutations frequently occur [4]. In practice, we also desire that two windows differing only by a few mutations are likely represented by the same *k*-mer in the sketch.

To quantify this notion of robustness, Edgar [4] proposes an alternative metric called *conservation*, which, given a sequence *S*, measures the expected number of *k*-mers that are persistently selected on some randomly mutated homologous sequence, divided by the sequence length. Edgar [4] argues that the minimizer method is context-dependent (i.e., the decision to sample any *k*-mer is made based on its neighboring *k*-mers), thus tends to have worse conservation scores compared to context-free methods that select *k*-mers based on their intrinsic content. This insight, which was later theoretically justified under reasonable assumptions by Shaw and Yu [19], motivated a seemingly new context-free sequence sketching method called *syncmers*, which samples every *k*-mer in which the lowest priority *s*-mer (*s* ≤ *k*) is found at some specific offset. Although both minimizers and syncmers employ the same concept of sampling based on a substring total ordering, a minimizer scheme using an *s*-mer ordering will directly report *s*-mers in its sketch, whereas a syncmer scheme parameterized by the same ordering will typically report longer *k*-mers. Because of this representation mismatch, all existing work [4, 19] unanimously chose to compare schemes that report similar-sized substrings regardless of their ordering choices, which results in an asymmetry of information among their sketches and prevents the derivation of any meaningful correspondence between minimizers and syncmers.

This shortcoming motivates a revision of the comparability notion for sketching methods, which subsequently leads to a rigorous theoretical understanding of their relationship. In particular, we seek to compare schemes that are parameterized by the same ordering and normalize the difference in their representations via selecting alternative modes of reporting that do not semantically change the sketches. This provides an invariant quantity among the compared methods, which allows us to derive formal bounds for their performance gaps in terms of density and conservation. Our theoretical result leads to a new sketching scheme called *masked minimizers*, which generalizes both minimizers and syncmers. We further introduce a new performance metric called the *generalized sketch score* (GSS) that combines both the traditional density metric and the conservation metric proposed by Edgar [4], resulting in the first unified protocol for comparing sequence sketches. Last, we propose an algorithm to optimize the masked minimizer scheme with respect to the proposed GSS metric. Our contributions are summarized as follows:

### Information-driven comparison of minimizers and syncmers

We propose a new notion of comparability for minimizers and syncmers, which focuses on contrasting schemes that use the same substring ordering to make sampling decisions. This allows us to explicitly bound the density and conservation gaps between comparable minimizers and syncmers. Our result further reveals that every syncmer sketch can be derived from its comparable minimizer sketch, which simultaneously has higher (better) conservation and higher (worse) density. This result is the first known correspondence between these sketching methods.

### Unifying minimizers and syncmers

We generalize this finding and introduce the concept of *masked minimizers* to unify both minimizers and syncmers. The masked minimizer scheme combines the standard minimizer sampling with an additional sub-sampling step via applying a binary mask to every window selection. Varying the mask configuration induces a spectrum of comparable schemes, including minimizers (i.e., all-ones mask) and various syncmer schemes (i.e., one-hot masks). This unification reveals a methodical approach to derive comparable sketching schemes via combining existing minimizer construction/optimization techniques [7, 5, 21, 22] with a mask optimization routine.

### Novel performance metric

We show that density and conservation are conflicting objectives that should not be considered in isolation and propose the GSS metric to robustly measure sequence sketching performance. Particularly, density is lower-bounded by conservation, thus increasing conservation will prevent improving density and vice versa. Optimizing for the ratio between conservation and density, or *relative conservation*, also leads to exploitative solutions that select unreasonably few *k*-mers and are ultimately meaningless as their sketches carry insufficient information about the sequence. The GSS metric addresses this issue by scaling the relative conservation metric with a regularization term that measures the *coverage* of the sketch with respect to the target sequence, hence penalizes trivial solutions.

### Conservation-aware optimization of masked minimizers

We propose a sequence-specific learning objective to optimize the masked minimizer scheme with respect to the GSS metric. Our objective function extends the DeepMinimizer loss function [7] with a secondary objective that minimizes the expected change in local ordering of *k*-mers under random mutations. Our approach consistently improves the GSS metric for every mask variant and outperform other known minimizer optimization approaches such as Miniception [21], PASHA [5] and DeepMinimizer [7]. We further conduct ablation studies to reveal practical insights about masked minimizers, such as the existence of trivial solutions, mask performance profiles and the ability to safeguard against a common minimizer pitfall.

Our masked minimizers approach generalizes and extends two broad classes of sequence sketching algorithms. This unifying perspective allows us to theoretically compare and optimize sketching schemes with heterogeneous mechanisms, which has not been achieved previously. Our method results in sketches that strike a balance between well-covering the target sequence and achieving favorable density-conservation trade-off, which will help to improve genomic analysis in applications that require sequence sketching.

## 2 Background and Notation

### Notation

Let *Σ* be an arbitrary alphabet. Given some parameter *k, a* (*w, k*)-window is defined as a substring of length 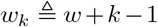, which contains exactly w overlapping *k*-mers. We further let *S* ∈ *Σ*^*L*+*k*–1^ be a string specified on *Σ* such that there are exactly *L* overlapping *k*-mers and *L_w_* ≜ *L* – *w* + 1 overlapping (*w, k*)-windows in *S*. Generally, we assume that *w* ≪ *L* and *k* ≪ *L*, as is typical in most practical settings. We use the notations 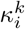 and 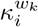 to respectively denote the *i*^th^ *k*-mer and the *i*^th^ (*w, k*)-window in *S*.

### *k*-mer sampling scheme

A *k*-mer sampling scheme 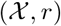 consists of: (1) a sampling function 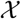 such that 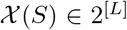 returns a set of locations in *S*; and (2) a one-to-one reporting function r that maps each sampled location in 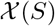 to a unique tuple of *k*-mer and *k*-mer location in *S*. The location distinguishes the same *k*-mer selected from different parts of the sequence. We note that the traditional definition of *k*-mer sampling schemes typically ignores the choice of *r* and assumes 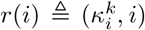, which directly reports the *k*-mer at the sampled location. The reporting function makes explicit the distinction between the sampling method to select locations (often the location of a distinguished *k*-mer) and the interpretation or use of these locations (e.g., the distinguished *k*-mer itself, a window containing this *k*-mer, the sequence between two consecutive selected locations [3]). This formalism unifies the behaviors of *k*-mer sampling schemes, including minimizers and syncmers. Some example reporting functions are demonstrated in Fig. 1. Lastly, the final reported sketch is given by the notation 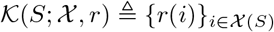.

**Fig. 1.**
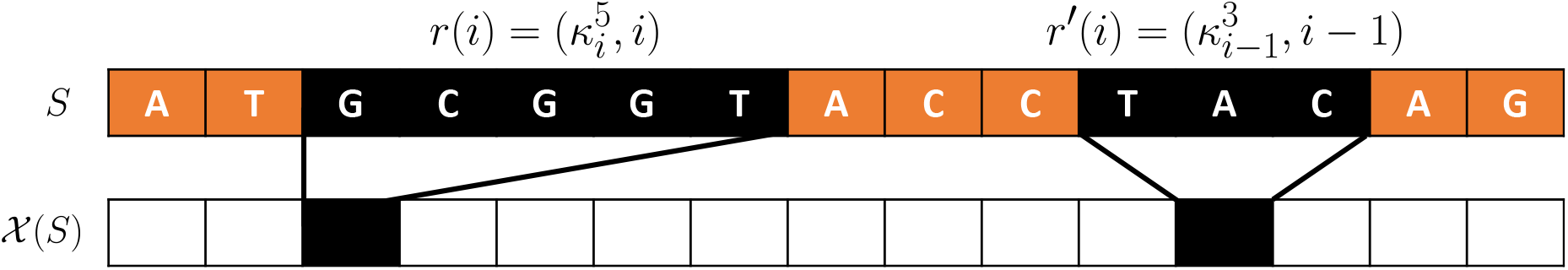
Illustrating two reporting functions *r*(*i*) = (*κ*^5^,*i*) and 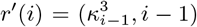. Black cells indicate the sampled locations in 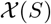 and the corresponding submers reported by each function. Depending on which function is used, the sampling function 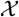 can be realized as a 5-mer sampling scheme 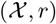 or a 3-mer sampling scheme 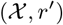.

### Minimizers

The sampling function of a minimizer scheme is characterized by a tuple of parameters (*w, k, π*), where *w* and *k* are defined above. Additionally, *π* is a total ordering over the set of all *k*-mers, which can be represented [7] as a scoring function 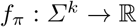, such that for every pair of *k*-mers *k, κ′* ∈ *Σ^k^*:

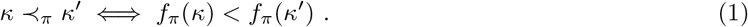

Here, 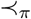 denotes the precedence of ordering in *π* and tie-breaking of equally scoring *k*-mers is determined by their order of appearance in the window. Let 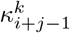 denote the *j*^th^ *k*-mer in the window 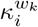, we define the selector function 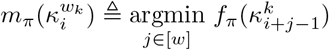, which returns the lowest scoring *k*-mer in some (*w, k*)-window of *S*. The minimizer sampling function is then given by iteratively applying *m_π_* to every window:

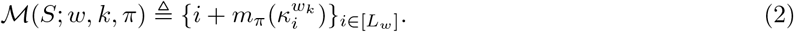

The minimizer scheme typically uses the identity reporting function 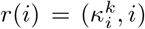 and is considered a *k*-mer sampling scheme. Schleimer et al. [18] used the concept of “charged windows” (a window selecting a different *k*-mer than the preceding window), effectively interpreting a (*w, k, π*)-minimizer scheme as a *w_k_*-mer (window) sampling scheme. The Miniception [21] minimizers scheme uses a similar idea of “charged context” to create an order between *k*-mers.

### Syncmers

While the minimizer scheme estimates the importance of a *k*-mer based on the relative ordering in its local neighborhood, the syncmer scheme instead uses the relative ordering of its constituent *s*-mers (i.e., *s* ≤ *k*). The closed-syncmer scheme is identical to a minimizer scheme reporting the charged windows. In the remainder of this paper we focus on the open-syncmer scheme, called simply syncmer for brevity.

The sampling function of a syncmer scheme is characterized by a tuple of parameters (*k, s, t, π*). The parameter *s* ≤ *k* indicates an implicit representation of *k*-mers as the cascaded sets of *k_s_* ≜ *k* + *s* – 1 overlapping *s*-mers. The total ordering *π* is instead defined on the set of all *s*-mers and can be likewise represented by the scoring function 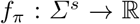. Let 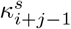 denote the *j*^th^ *s*-mer in 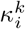, we define a similar selector function 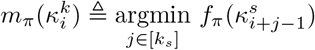, which returns the lowest ranking *s*-mer in 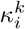, for some 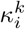 in *S*. Unlike minimizers, however, syncmers only sample a location *i* if the lowest ranking *s*-mer in 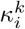 is at the *t*^th^ position, i.e., 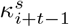. The syncmer sampling function is given by iteratively applying this to every *k*-mer:

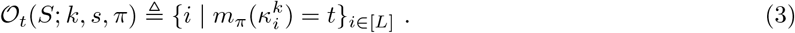

The syncmer scheme also commonly adopts the identity reporting function 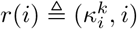. However, while the minimizer scheme guarantees that every (*w, k*)-window contains at least one sampled location, the syncmer scheme does not. Lastly, as our analyses tend to fix the settings of *w, k, s* and *π*, we typically suppress these parameters and shorten the above notations to 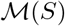 and 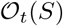, unless otherwise necessary.

### Density metric

Let 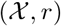 be an arbitrary sampling scheme where *r* reports *k*-mers, the density metric measures the size of the sketch 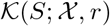 relative to the number of *k*-mer in *S* (lower is better):

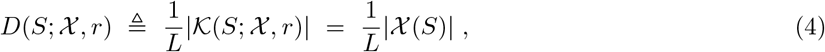

where the second equality is due to r being a one-to-one mapping.

### Conservation metric

The conservation metric [4] measures the expected number of sketch items that are present in both 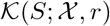 and 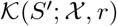 relative to the number of *k*-mers in *S* (higher is better), where *S*′ is a randomly mutated copy of *S* following some arbitrary distribution *p_S_*:

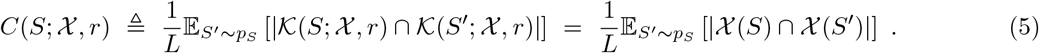

## 3 Comparing and Unifying Sketching Schemes

### 3.1 Revisiting performance metrics

Let 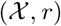 be an arbitrary sampling scheme. As the set of conserved locations 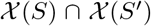 is always a subset of 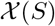 regardless of the mutations in *S*′, we have 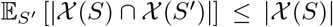, and therefore 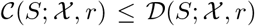 for all *S*. Improving (lowering) density thus places an upper-bound on how much conservation can be improved and vice versa. This conflicting nature between the two objectives implies that neither should be considered independently of one another. To account for this trade-off, we propose a new metric called the *generalized sketch score* (GSS), which is defined as follows:

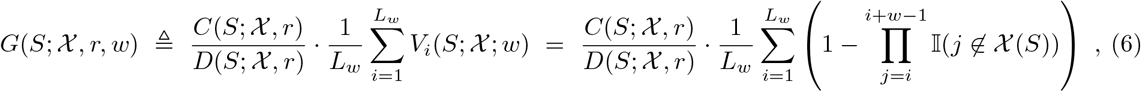

where 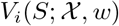 is the indicator variable of the event 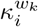 overlaps at least one sampled location in 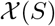.

Explicitly, our metric consists of two components. The first component, 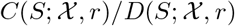, evaluates the *relative conservation* and captures the inherent trade-off between the density and conservation metrics. Interestingly, this term also corresponds to measuring the number of conserved locations in 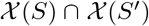 relative to the number of sampled locations in the original sketch 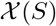:

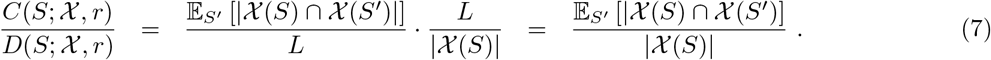

As a metric, this term alone is, however, vulnerable to a simple exploit that trivially maximizes it. To see this, we first note that around each sampled location, there is a finite-length substring (i.e., context) in which a mutation can possibly alter the sampling outcome. With respect to the default reporting function, the context of a minimizer-sampled location is the union of all windows that contain it. The context of a syncmer-sampled location is the *k*-mer at the same position. Independent of the random mutations, the portion of the sequence in which mutations might induce an effect on the relative conservation metric is therefore bounded by the number of selected locations in 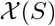. Therefore, the fewer locations sampled by 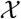 means a smaller probability for any mutation to occur in this conservation-sensitive portion of the sequence.

Naturally, if the sketch picks an unreasonably small number of locations, it is likely that the relative conservation term will be near perfect (i.e., close to 1). One such scenario could theoretically occur with the syncmer setting, when 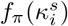 is constructed such that the lowest scoring *s*-mer in every *k*-mer is always found within the first *t* – 1 positions. When this happens, a large portion of the sequence is not represented by any *k*-mer, thus resulting in a meaningless sketch. We demonstrate that such a sampling behavior can be found via optimization in Section 5. To prevent this sub-optimal outcome, our second term 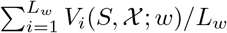 measures the *coverage* of the sketch, or the fraction of (*w, k*)-windows that contains at least one sampled *k*-mer. When very few *k*-mers are selected, the resulting low coverage will apply a discount on the high relative conservation term, hence will discourage these trivial solutions.

### 3.2 Comparable minimizers and syncmers

Minimizer and syncmer schemes with similar *k* are typically deemed comparable [4, 19] since they both adopt the *k*-mer reporting function 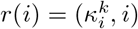. This notion of comparability, however, does not facilitate a theoretical analysis of their performance differences (i.e., in terms of density and conservation metrics), as different information bases are used to enact the sampling decisions of a length-k minimizer and syncmer. In particular, (*w, k*)-minimizer schemes use total *k*-mer orderings to perform sampling, whereas (*k, s, t*)-syncmer schemes use total *s*-mer orderings, with *s* ≤ *k*. We note that these bases are only comparable when setting *s* = *k*, but doing so results in trivial syncmer schemes that selects every *k*-mer in *S*.

To correct this asymmetry of information, we propose to compare *π*-comparable minimizers and syncmers, which are sketching schemes that employ the same total ordering π. For example, the (*w, k, π*)-minimizer and the (*w_k_, k, t, π*)-syncmer schemes with *w_k_* ≜ *w* + *k* – 1 and *t* ≤ *w*, which respectively report *k*-mers and *w_k_*-mers, are *π*-comparable. To work around their difference in representation, we further replace the default *w_k_*-mer reporting function 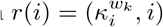 of the above syncmer scheme with the *k*-syncmer reporting function 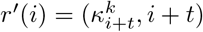. As both reporting functions are one-to-one mappings of the selected locations, this substitution results in a semantically equivalent *shifted* syncmer scheme that reports all lowest ranking *k*-mers that are at the *t*^th^ position in their respective w_k_-mers, rather than the w_k_-mers themselves.

This translation of the reporting function aligns our proposed comparison with the traditional perspective of comparability and presents an invariant substring ranking behavior among the compared schemes, which subsequently allows us to derive explicit bounds on their density and conservation gaps. In particular, Proposition 1 proves that the sketch of any syncmer is a subset of its *π*-comparable minimizer sketch. Corollary 1 and Corollary 2 further show that the density and conservation of any syncmer on a specific sequence S are respectively upper-bounded and almost surely upper-bounded by that of its *π*-comparable minimizer, thus establishing the first theoretical correspondence between *π*-comparable schemes.

#### Proposition 1.

*Given* 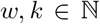 *and a total ordering π defined on the set of all *k*-mers, we let* (*w, k, π*) *and the reporting function* 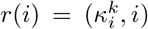 *define a minimizer scheme* 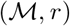. *Further let* (*w_k_, k, t, π*) *and* 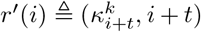 *define a shifted syncmer scheme* 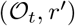, *such that t* ≤ *w*. Then, for all S ∈ *Σ*^*L+k*–1^, *we have* 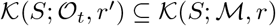.

*Proof.* We first note that 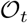 will not sample any location *i* such that 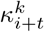, or the *t*^th^ *k*-mer in 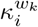, does not exist, hence *r*′ is well-defined. Then, by definition of *r* and *r*′, it suffices to show that 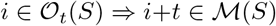 for all *i* ∈ [*L_w_*]. As both schemes are parameterized by the same ordering *π*, we can express their respective sets of sampled locations using the same selector function *m_π_*:

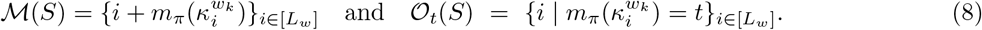

Therefore, for all *i* ∈ [*L_w_*], we have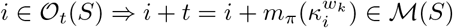.

#### Corollary 1 (Density gap of *π*-comparable schemes).

*Let* (*w_k_, k, t, π, r′*) *and* (*w,k,π,r*) *define a pair of π-comparable shifted syncmer* 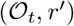 *and minimizer* 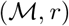 *schemes as described in Proposition 1, then for all S* ∈ *∑*^*L*+*k*–1^, *we have* 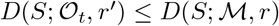.

*Proof.* This follows directly from Proposition 1, which establishes that 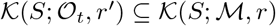. Thus, we have 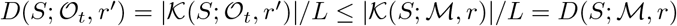.

#### Corollary 2 (Conservation gap of *π*-comparable schemes).

*Let* (*w_k_, k, t, π, r′*) *and* (*w,k,π,r*) *define a pair of π-comparable shifted syncmer* 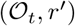 *and minimizer* 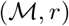 *schemes as described in Proposition 1, then for all S* ∈ *∑*^*L*+*k*–1^, *we have* 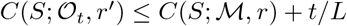.

*Proof.* Our proof is given in Appendix A.

We note that since *t* ≤ *k* ≪ *L* in most practical applications, Corollary 2 almost surely corresponds to an exact upper bound. Finally, we state our main result in Theorem 1 below, which establishes the first theoretical correspondence between minimizers and syncmers. That is, for every syncmer, there exists a corresponding minimizer scheme that simultaneously yields higher (worse) density and higher (better) conservation.

#### Theorem 1 (*π*-comparability defines a correspondence between minimizers and syncmers).

*For every syncmer scheme* 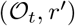, *there exists a comparable minimizer scheme* 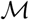 *whose reporting function r is translated from r′ with an offset t, such that the following bounds hold for every S* ∈ *Σ*^*L*+*k*–1^:

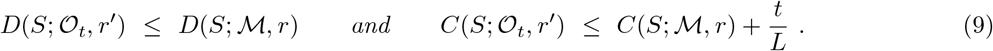

*Proof.* The proof of Theorem 1 follows directly from Corollary 1 and Corollary 2.

### 3.3 Unifying comparable schemes with masked minimizers

Proposition 1 establishes that 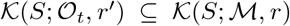 for any *π*-comparable pair of schemes 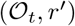 and 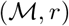. Additionally, it follows from the definitions of 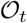 and 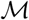 that, for all 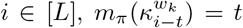 simultaneously implies i ∈ 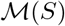 and 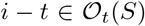. Thus, 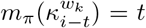 suffices as a sub-sampling rule to recover 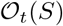 given 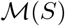, or likewise to recover 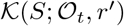 given 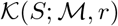. We write this rule as:

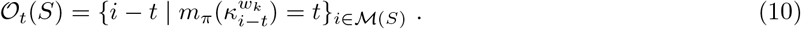

Let the output of the selector function *m_π_* be equivalently represented as a *w*-dimensional one-hot vector 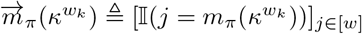, where *κ^w_k_^* is some arbitrary (*w, k*)-window in *S*. The above sub-sampling rule can then be rewritten as the point-wise multiplication between 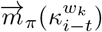 and the *w*-dimensional one-hot vector 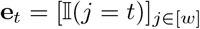, whose 1-entry is at the *t*^th^ position:

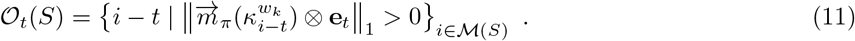

In the above formulation, the positive 1-norm condition checks if any position remains selected after filtering with **e**_*t*_ (in which case, it must be the *t*^th^ position). Interestingly, this sub-sampling rule can be generalized by replacing **e**_*t*_ with any arbitrary w-dimensional binary mask *v* ∈ {0,1}^*w*^. For example, setting *v* = **1**_*w*_ trivially recovers the standard minimizer scheme (without applying the offset *t*). Given a set of minimizer-sampled locations 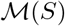, varying v yields a total of 2^*w*^ *π*-comparable schemes, leading to a unifying method called *masked minimizers.* We state its definition below.

#### Masked minimizers

The sampling function of a masked minimizer scheme is characterized by a tuple of parameters (*w, k, π, v*), where *w, k, π* correspond to standard minimizer parameters, and *v* ∈ {0,1}^*w*^ is a *w*-dimensional binary vector. The masked minimizer sampling function is given by:

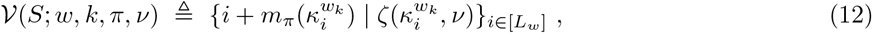

where 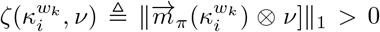 denotes the event that the selection at the *i*^th^ window remains sampled after applying the sub-sampling mask. We note that this definition does not factor in the offset t to recover exactly the locations sampled by a comparable syncmers. We can, however, specify the reporting function 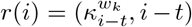 to recover 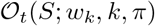 from 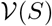. Thus, every syncmer and minimizer scheme can be written as a masked minimizer.

## 4 Optimizing masked minimizers

Following the standard setting of minimizer optimization, we fix *w, k* for the masked minimizer scheme and optimize for *π, v*. We address this objective via a bi-level optimization routine, which first optimizes for π, then performs greedy search for the optimal sub-sampling mask *v*, which starts with the minimizer mask **1**_*w*_ and iteratively zeroes out a position that yields the best performance improvement. The search terminates when no further improvement can be obtained. In principle, any existing optimizer for minimizer [5, 21, 22, 7] can be used in the first step. A full review of these methods is given in Appendix B. However, we note that all existing algorithms to optimize minimizers do not account for the conservation component that is reflected in the GSS metric. To address this, we adapt the DeepMinimizer loss function [7] with a secondary objective that minimizes the expected change in *k*-mer score assignment with respect to random mutations, hence encouraging high conservation.

The DeepMinimizer method [7] is a gradient-based reformulation of the intractable permutation learning task (i.e., learning *π*), which employs a pair of dueling neural networks to obtain a scoring function that induces low density on *S*. The PriorityNet 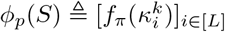 implicitly parameterizes *f_π_*, applies it on every *k*-mer in *S* and returns a priority score vector in ℝ^*L*^. The TemplateNet *ϕ_t_*(*S*) ≜ [*g*(*i*)]_*i*∈[*L*]_ applies a sequence-agnostic scoring function 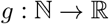 to every location in ℝ^*L*^. The parameterizations of *φ_p_* and *φ_t_* respectively ensure the validity of a minimizer scheme and the ideal low density property, but neither can initially achieve both. Hoang et al. [7] proposes to minimize a specially constructed distance *Δ*(*ϕ_p_*(*S*), *ϕ_t_*(*S*)) (which behaves similarly to a weighted distance) between their outputs in order to obtain a valid minimizer scheme that likely has low density on *S*. The details of *Δ, ϕ_p_* and *ϕ_t_* are given in Appendix C. Building upon this insight, our adapted loss function to optimize GSS is given as follows:

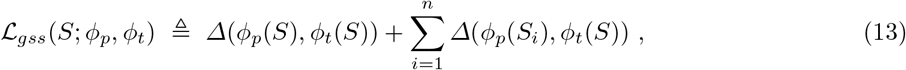

where *S*_1_, *S*_2_,…, *S_n_* denote *n* randomly sampled mutations of S. The first term on the right hand side is exactly the DeepMinimizer loss [7], which was designed to optimize for density. The second term minimizes the expected distance between each priority vector *ϕ_p_*(*S_i_*) for mutated sequence *S_i_* and the original template vector *ϕ_t_*(*S*). Intuitively, when this term is small, we expect {*ϕ_p_*(*S_i_*)}_*i*∈[*n*]_ to be concentrated around the template *ϕ_t_*(*S*). As *ϕ_t_*(*S*) is close to *ϕ_p_*(*S*) via minimizing the first term, this optimality implies that the score assignment *ϕ_p_*(*S*) is likely to be conserved under random mutations.

## 5 Empirical results

Our experiments aim to investigate the following: (1) Are density and conservation adversarially related? (2) How do *π*-comparable schemes perform relative to one another under the proposed metric GSS? (3) Can mask optimization improve the overall performance of both minimizer and syncmer? Through these demonstrations, we confirm our theoretical understanding of various sketching metrics and the relationship among *π*-comparable schemes, as well as demonstrate the efficiency of our proposed optimization method. We further conduct ablation studies to investigate several interesting phenomena related to masked minimizers, such as the ability to exploit the relative ordering metric (as mentioned in Section 3.1); the performance of all masks, which reveals insight on setting a good default mask; and the ability to prevent repeated sampling in a homopolymer-rich sequence, which has previously been a unique advantage of the syncmer method.

### Experimentation details

We compare the following methods to construct the *k*-mer ordering: (1) random ordering; (2) training with different variants of the DeepMinimizer loss [7]; (3) constructing the ordering from Miniception [21]; and (4) PASHA universal hitting sets [5]. Performance is demonstrated on the following mask parameters: (1) the minimizer mask *v* = **1**_*w*_; (2) the syncmer mask with offset *t* = *w*/2 (suggested by Shaw and Yu [19]) meaning *v* = **e**_*w*/2_; (3) its complement *v* = **1**_*w*_ — **e**_*w*/2_; and (4) the optimized mask *v*_*_ found by the heuristic search described above. We respectively label the sketches of these masked minimizers as 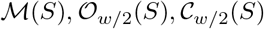 and 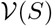. All experiments are trained on human chromosome 1 (labelled Chr1); the centromere region of human chromosome X (labelled ChrXC); and several bacterial genomes that were previously used in Edgar [4] (labelled Btr1, Btr2, Btr3 and Btr4). The details of these sequences are given in Appendix D. We compute all gradient-based loss functions per batch of sampled subsequences since it is not possible to fit the entire sequence on GPU memory. Other implementation details are given in Appendix E. Our implementation can be found at https://github.com/hqminh/maskedminimizer.

### The adversarial relationship of density and conservation

This experiment demonstrates that density and conservation are conflicting objectives for *π*-comparable schemes, thus confirming our analysis in Section 3.1. First, we train masked minimizers for *w* = 7, *k* = 15 and various binary masks *v* using three different loss functions: (1) the DeepMinimizer loss, which only optimizes for density and is labelled as 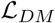; (2) only the conservation term in Eq. (13), which optimizes for conservation and is labelled as 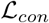; and (3) the combined masked minimizer loss, which is given as Eq. (13) and labelled as 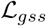.

Fig. 2 plots the density, conservation, coverage, relative conservation and GSS metrics as these masked minimizer schemes are trained with each loss function over 300 epochs on Btr1. As predicted in Section 3.1, we observe that the density metric is consistently greater than the conservation metric in all experiments. Training with any variant of the DeepMinimizer 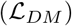 loss generally lowers density for minimizer 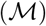 and complement 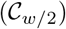 schemes, but also decreases their conservation. On the contrary, the conservation of syncmers 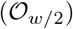 increases with training at the expense of raising its density. This is most likely because the sampling behavior of syncmers is not compatible with the above variants of 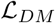, which was originally designed for minimizers; whereas the complement mask behaves almost identically to the minimizer mask due to its mild sub-sampling.

**Fig. 2.**
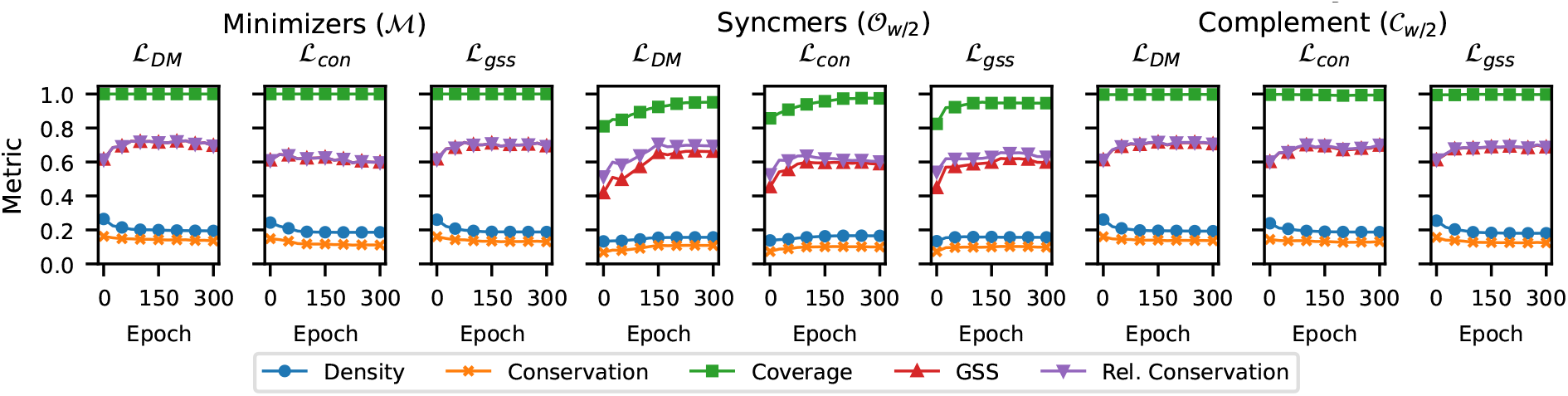
Comparing density, conservation, relative conservation and GSS vs. number of training epochs using difference training losses and masks on the bacterial genome Btr1.

### The effectiveness of training masked minimizers

This experiment demonstrates that our proposed loss function 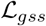 learns robustly and improves GSS in various settings of *w, k* and different masks *v*. Fig. 3 plots the GSS of the masked minimizers 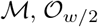 and 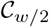 over 600 training epochs in two settings: (1) *w* = 15 and *k* ∈ [25, 40, 55, 70]; (2) *k* =15 and *w* ∈ [25, 40, 55, 70]. This experiment is repeated on two sequences, ChrXC and Chr1. All experiments show that GSS steadily increases over 600 training epochs by 1.5 to 5 times that of their initial random weights. We observe that the performance of minimizers 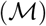 is highly similar to the complement scheme 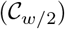, except for (*w, k*) = (15, 40) with Chr1 and (*w, k*) = (15, 25), (40,15) with ChrXC. Both these masks outperform syncmers 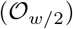 in most settings, which corroborates the observation in the previous experiment. Appendix F shows the individual effects of training on the conservation and density metrics for the experiments in Fig. 3(a), thus confirming our analysis in Section 3.2.

**Fig. 3.**
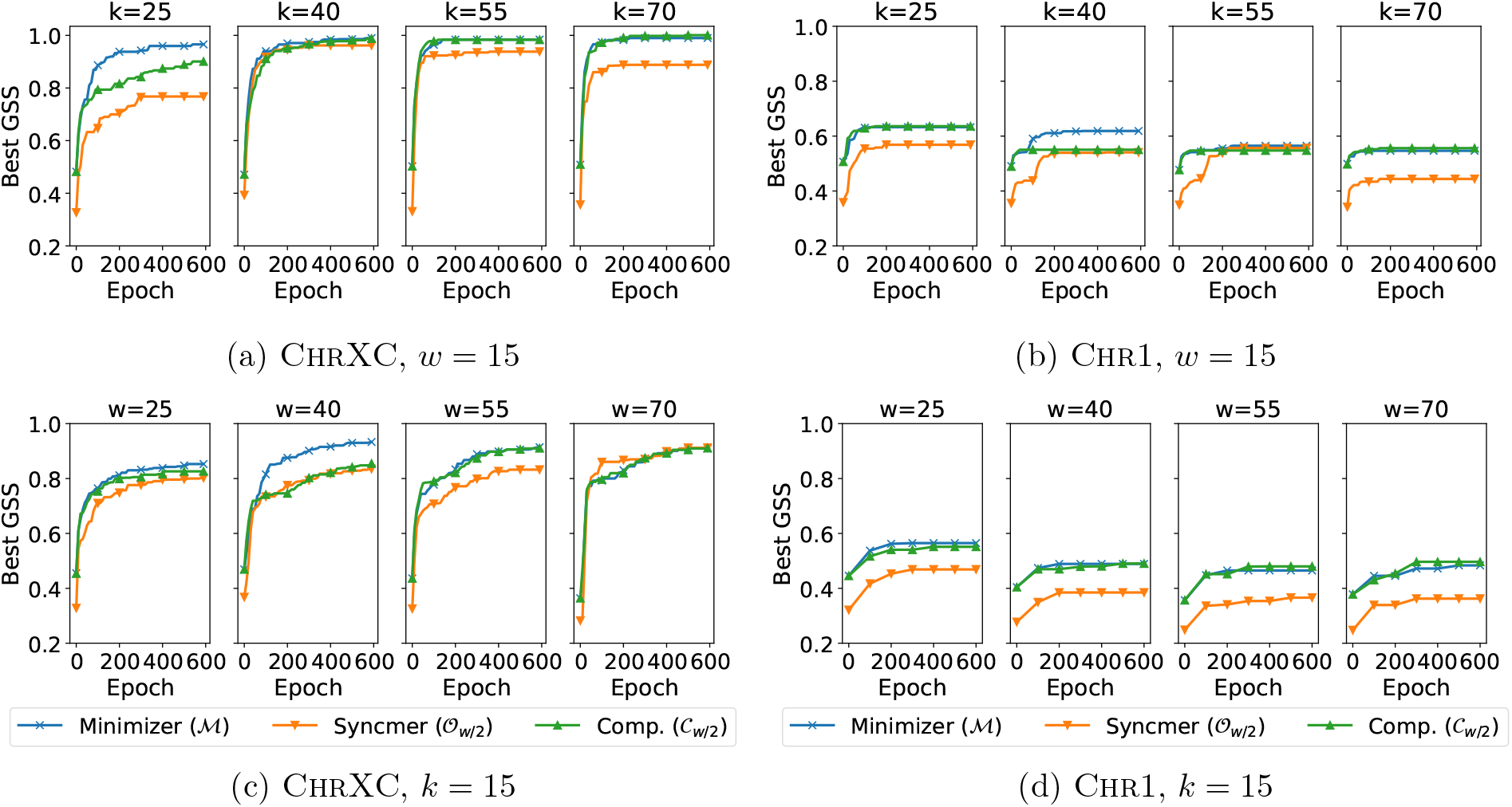
Comparing GSS of different, masked minimizers vs. number of training epochs on ChrXC and Chr1.

### Comparing GSS of compatible schemes with different training losses and masks

This experiment compares the performance of various masked minimizers whose orderings are randomized; trained with 3 different DeepMinimizer-based losses (i.e., 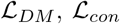 and 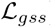); or constructed using other optimization techniques such as Miniception] and PASHA [5]. We compute the GSS on all combinations of *w* ∈ {10, 15, 20} and *k* ∈ {10, 15}. Table 1 summarizes the result of this study on ChrXC. Overall, the masked minimizer loss function 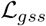 performs the best, having achieved the highest GSS in 4 over 6 settings of *w, k.* Across 18 experiments (i.e., crossing 6 settings of (*w, k*) and 3 loss functions), the best GSS is achieved by the minimizer mask on 6 experiments, the syncmer mask on 1 experiment, and the complement mask on 4 experiments. In 10 out of these 11 experiments, with (15, 15, 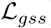) being the exception, the optimized mask *v*_≠_ achieves the same best GSS, which is reasonable because our greedy mask pruning algorithm does not guarantee finding the optimal mask. In the remaining 7 experiments, *v*_≠_ outperforms all handcrafted masks, thus confirming the need to conduct this optimization step.

**Table 1.** Comparing GSS of different masked minimizers with 3 different training losses 6 settings of (*w,k*) on ChrXC. The best GSS observed for each combination of (*w, k*) and loss function is given in **bold**.

| $(w, k)$ | Conservation Loss $\mathcal{L}_{con}$ | | | | DeepMinimizer Loss $\mathcal{L}_{DM}$ | | | | Masked Minimizer Loss $\mathcal{L}_{gss}$ | | | |
| --- | --- | --- | --- | --- | --- | --- | --- | --- | --- | --- | --- | --- |
| | $\mathcal{M}$ | $\mathcal{O}_{w/2}$ | $\mathcal{C}_{w/2}$ | $\mathcal{V}$ | $\mathcal{M}$ | $\mathcal{O}_{w/2}$ | $\mathcal{C}_{w/2}$ | $\mathcal{V}$ | $\mathcal{M}$ | $\mathcal{O}_{w/2}$ | $\mathcal{C}_{w/2}$ | $\mathcal{V}$ |
| 10, 10 | **0.701** | 0.656 | 0.685 | **0.701** | 0.716 | 0.529 | 0.722 | **0.745** | **0.754** | 0.601 | 0.753 | **0.754** |
| 10, 15 | **0.706** | 0.650 | 0.696 | **0.706** | 0.768 | 0.656 | **0.779** | **0.779** | 0.719 | 0.566 | 0.704 | **0.810** |
| 15, 10 | 0.693 | 0.621 | 0.680 | **0.713** | 0.690 | 0.613 | 0.726 | **0.765** | 0.683 | 0.597 | 0.701 | **0.750** |
| 15, 15 | 0.715 | 0.667 | 0.710 | **0.892** | 0.813 | 0.746 | **0.828** | **0.828** | 0.812 | **0.823** | 0.817 | 0.817 |
| 20, 10 | **0.682** | 0.614 | 0.679 | **0.682** | 0.677 | 0.608 | 0.673 | **0.690** | **0.721** | 0.595 | 0.690 | **0.721** |
| 20, 15 | 0.747 | 0.716 | **0.816** | **0.816** | 0.829 | 0.714 | **0.845** | **0.845** | **0.898** | 0.794 | 0.894 | **0.898** |

We repeat the same experiment for non-gradient optimization methods, including PASHA [5], Miniception[21] and the random ordering baseline. We summarize their GSS performance in Table 2. Among these methods, PASHA obtains the best GSS in 4 over 6 (*w, k*) settings, whereas Miniception obtains the best GSS in the other 2 settings. While these methods outperform the random ordering baseline as expected, their performance is generally weaker than the gradient-based methods above. Similar to the previous experiment, we also observe that *v*_≠_ achieves the best on 17 over 18 settings with 10 being clear improvements over handcrafted masks. More interestingly, the syncmer mask obtains a GSS of 0.0 with Miniception in many settings of (*w, k*), which suggests that none of the sampled locations is found at the *w/2* offset. Although the cause of this is unclear, this phenomenon confirms the necessity of finding an optimal mask.

**Table 2.** Comparing GSS of different masked minimizers with 3 different discrete construction methods and 6 settings of (*w, k*) on ChrXC. The best GSS observed for each combination of (*w, k*) and construction method is given in **bold**.

| $(w, k)$ | MINICEPTION UHS | | | | PASHA UHS | | | | Random Ordering | | | |
| --- | --- | --- | --- | --- | --- | --- | --- | --- | --- | --- | --- | --- |
| | $\mathcal{M}$ | $\mathcal{O}_{w/2}$ | $\mathcal{C}_{w/2}$ | $\mathcal{V}$ | $\mathcal{M}$ | $\mathcal{O}_{w/2}$ | $\mathcal{C}_{w/2}$ | $\mathcal{V}$ | $\mathcal{M}$ | $\mathcal{O}_{w/2}$ | $\mathcal{C}_{w/2}$ | $\mathcal{V}$ |
| 10, 10 | 0.574 | 0.000 | 0.551 | **0.582** | **0.629** | 0.438 | 0.601 | **0.629** | 0.265 | 0.210 | 0.287 | **0.288** |
| 10, 15 | 0.473 | 0.251 | 0.487 | **0.501** | 0.756 | 0.192 | **0.769** | **0.769** | 0.222 | 0.079 | 0.262 | **0.265** |
| 15, 10 | **0.557** | 0.000 | 0.576 | **0.557** | 0.510 | 0.444 | **0.589** | **0.589** | 0.271 | **0.276** | 0.253 | 0.271 |
| 15, 15 | 0.435 | 0.369 | 0.471 | **0.509** | 0.521 | 0.302 | 0.558 | **0.632** | **0.172** | 0.119 | 0.140 | **0.172** |
| 20, 10 | **0.604** | 0.000 | 0.489 | **0.604** | 0.435 | 0.307 | 0.551 | **0.553** | 0.189 | 0.202 | 0.239 | **0.247** |
| 20, 15 | 0.392 | 0.000 | 0.435 | **0.476** | 0.329 | 0.315 | 0.390 | **0.399** | **0.132** | 0.095 | 0.128 | **0.132** |

### Exploiting the relative density metric

This experiment demonstrates the exploitative behavior mentioned in Section 3.1 via a special loss function 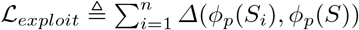, which differs from 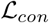 by swapping the template *ϕ_t_* (*S*) in each pairwise *Δ*-distance term with *ϕ_p_*(*S*). The purpose of this substitution is to isolate any training signal for density (which is implicitly encoded in the template) and directly prioritize minimizing relative conservation. As minimizers schemes must select one position per (*w, k*)-window by construction, they do not suffer from this exploit. We thus train only the syncmer mask 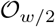 on a random sequence with *L* = 1000, using 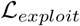 with *w* = 10 and *k* = 15.

Fig. 4 (left) plots the relative conservation and coverage metrics obtained over 1000 epochs with different number of sampled mutations *n* ∈ {1, 5, 10, 20} per training epoch. We observe that 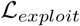 consistently improves relative conservation as expected. When n = 20, the optimizer finds the exploit mentioned in Section 3 after 1000 epochs, which causes both metrics to reach 0. The resulting sketch selects no *k*-mers (i.e., 0 coverage) and is trivially conserved when mutations are introduced (i.e., infinity conservation, manually set to 0). Fig. 4 (right) plots the number of segments with monotonically increasing/decreasing priority scores at each segment length. The longest segment with monotonically decreasing priority score is 4, which is smaller than the syncmer offset *t* = *w*/2 = 5. This implies that all lowest scoring *k*-mers are found within the first *t* – 1 positions of their respective windows and none are sub-sampled into the syncmer sketch. Appendix F repeats this experiment for *t* = 6, 7, 8, 9 and yields the same pattern with their discovered exploits, thus confirming the existence of our theoretical scenario in Section 3.1 and the necessity of the GSS metric.

**Fig. 4.**
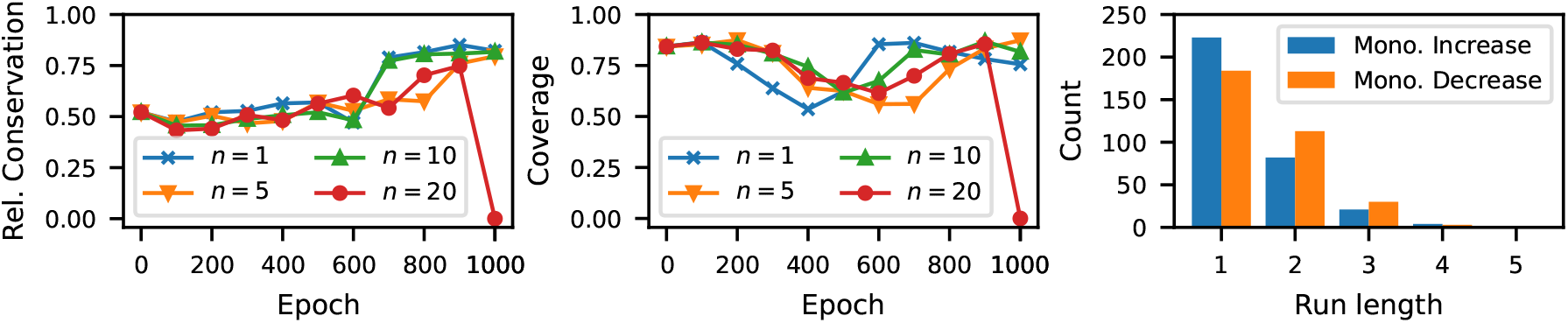
Left: relative conservation and coverage of 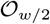 trained on a random sequence using 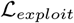 with varying no. sampled mutations *n*; Right: no. segments with monotonically changing priority scores at each segment length.

### The minimizer mask is a good default configuration

Fig. 5 (left) shows the scatter plot of all 2^*w*^ masked minimizers trained on Btr4 using 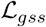 with *w* = 10 and *k* = 15, grouped by the number of 1-entries in their masks. Similar experiments on Btr1, Btr2 and Btr3 are deferred to Appendix F. We observe that the average GSS increases with the number of 1-entries in all experiments, which implies that the minimizer mask is a good default choice. However, we also observe that there exist specific masks that perform better than the minimizer mask, thus justifying mask optimization in certain applications.

**Fig. 5.**
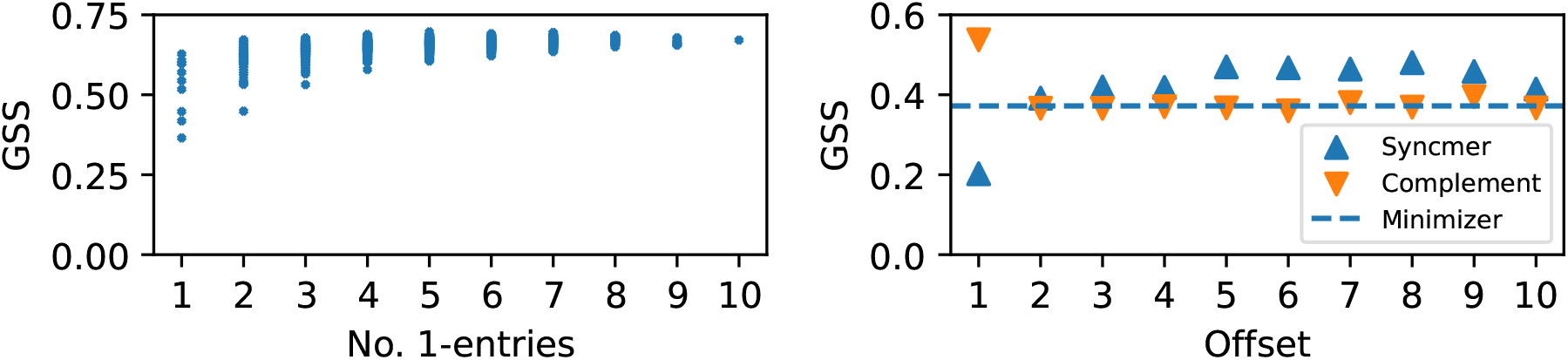
Left: GSS vs. number of 1-entries of all mask minimizers trained on the bacterial genome Btr4; Right: GSS vs. offset positions of syncmer and complement masks on a synthetic sequence with high homopolymer content.

### Preventing repeated sampling in homopolymer-rich sequences

One advantage of syncmers with *t* > 1 is the ability to avoid repeated sampling of identical *k*-mers in homopolymer substrings (i.e., substrings with repeated submer patterns) [4]. To confirm this, Fig. 5 (right) plots the GSS of all syncmer masks and their complements on a synthetic sequence with *L* = 100000 and 0.2% homopolymer content. The dotted line shows the GSS of the minimizer mask, which expectedly performs worse than most syncmers (except for *t* = 1) due to the repeated sampling pitfall. Because of the left-most tie breaking rule, every masked minimizers with a 1 at the left most position of the mask (which includes minimizers, syncmers with *t*=1 and complement with *t* ≠ 1) suffer from the pitfall of having a high density. This is indeed reflected in their similar GSS to the minimizer mask. On the contrary, we observe that the complement mask 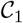 (i.e., 1^st^ position is pruned) achieves the best GSS of 0.56. This is because it avoids the repeated sampling pitfall in the same way any syncmer mask with *t* > 1 does, but otherwise performs like a minimizer scheme and does not suffer from the low coverage of syncmers.

## 6 Conclusion

We introduced: (1) a *π*-comparability notion for sequence sketching schemes; (2) a robust metric called GSS that can be used to contrast the performance of comparable schemes; and (3) a novel framework called masked minimizers that unifies comparable schemes, thus revealing a methodical approach to optimize them. We propose a practical optimization technique for masked minimizers and demonstrate that our method finds sketches with better density-conservation ratio than other sketch construction methods while maintaining high coverage of the sequence on various practical settings. Our empirical results confirm our theory and reveal important insights regarding sequence sketching. This research thus opens up new directions for systematic construction of sequence sketches that improve genomic analysis. Our future work will investigate better approaches to the masked minimizer optimization problem, as the current iteration is still crude and somewhat inefficient, due to the repeated computation of mutation priority scores in the loss function.

## Acknowledgements

This work was supported in part by the US National Science Foundation [DBI-1937540], the US National Institutes of Health [R01HG012470] and by the generosity of Eric and Wendy Schmidt by recommendation of the Schmidt Futures program. Conflict of Interest: C.K. is a co-founder of Ocean Genomics, Inc.

## A Proof of Corollary 2

### Corollary 2 (Conservation gap of *π*-comparable schemes).

*Let* (*w_k_, k, t, π, r′*) *and* (*w,k,π,r*) *define a pair of π-comparable shifted syncmer* 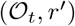 *and minimizer* 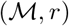 *schemes as described in Proposition 1, then for all S* ∈ *∑*^*L*+*k*–1^, *we have* 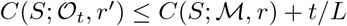.

*Proof.* Fixing a mutation *S*′ and let 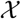 be an arbitrary location sampling function, we additionally define the indicator variable 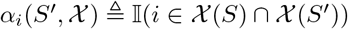 of the event that location *i* is preserved across 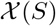 and 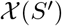. We first give the following bound of 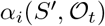 in terms of 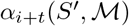:

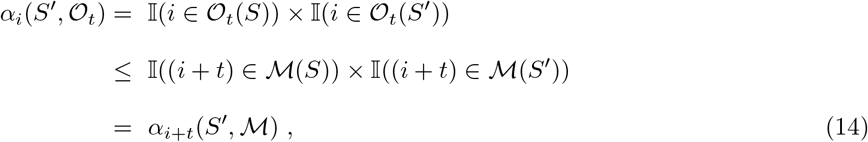

where the inequality follows from Proposition 1, which establishes that 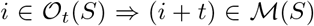. As *r* and *r*′ are both one-to-one mappings, we can rewrite the conservation difference between 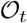 and 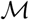 as:

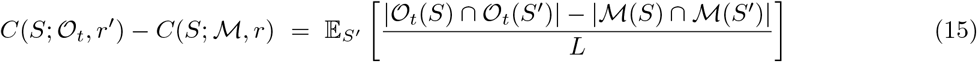

Expressing the cardinality of each intersection set above as the sum of its indicator variables gives:

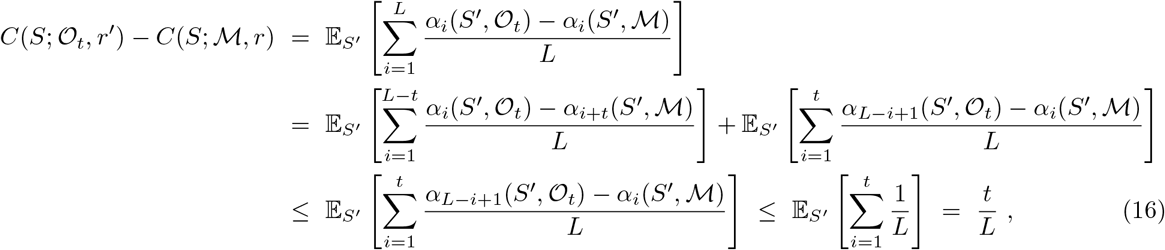

where the first inequality in the above derivation follows from the bound in Eq. (14), and the second inequality is due to the fact that each *α* term is either 0 or 1. Rearranging the above result concludes our proof.

## B Minimizer optimization and selection methods

### Heuristic methods

A random ordering is the most commonly adopted selection method for minimizer. The expected density of a random ordering given a window length *w* is 2/*w*. Beyond this scheme, several other methods rank *k*-mers based on their frequencies in the target sequence [1, 10] or sequentially remove *k*-mers from some arbitrarily constructed UHS [2].

### Priority set methods

Minimizer selection schemes with expected performance guarantees are based on the theory of universal hitting sets (UHS) [13, 16]. A (*w, k*)-UHS is defined as a set of *k*-mers such that every window of length *w* (from any possible sequence) contains at least one of its elements. A UHS can be thought of as a less fine-grained scoring function *f_π_*, which outputs only two score values, one for every *k*-mer in the set, and a larger score for every other *k*-mer (i.e., lower priorities). A more compact UHS has been shown to correlate with lower density, hence most UHS-based methods such as Miniception and PASHA are representative approaches that focus on minimizing the size of UHS. Zheng et al. alternatively adopts the concept of a polar set, whose elements are sufficiently far apart on a specific target sequence [22], thus extending this optimization paradigm to the sequence-specific setting. The polar set optimization objective, however, is NP-hard and currently approximated by a greedy construction [22].

### Gradient-based method

Hoang et al. recently propose the first relaxation for the discrete ordering optimization underlying the minimizer selection problem [7]. This involves two dueling networks: the PriorityNet focuses on constructing valid minimizer scheme (i.e., a total ordering can be reconstructed given the network), whereas the TemplateNet focuses on finding a scoring function that has few local optima (i.e., thus implying low density). The proposed loss function minimizes a special distance metric *Δ* between the output of these networks (given *S*), thus inducing a consensus solution that is valid and has low density on *S*. The architectures of these networks and the distance function *Δ* are given in Appendix C.

## C Parameterization of the DeepMinimizer network

### Input representation

Let *B* denote a batch of *b* subsequences of length ℓ; *B_i_* denote the *i*^th^ subsequence in the batch; *B_i,j_* denote its *j*^th^ character; 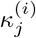 denote its *j*^th^ *k*-mer; and 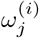 denote its *j*^th^ (*w, k*)-window.

### PriorityNet

The PriorityNet *ϕ_p_* takes as input the multi-hot representation of *B*, which is a tensor *T* ∈ {0,1}^*b*×ℓ×|*∑*|^, where *T_i,j_* = **e**_*c*_ if and only if *B_i,j_* is indexed *c* in *Σ*. The output of *ϕ_p_* is a priority score matrix *P* ∈ ℓ^*b*×*ℓ*^ with the semantic 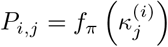. The PriorityNet *ϕ_p_* is parameterized by a 3-layered convolutional neural network. The first layer has filter size *k* and subsequent layers have filter sizes 1 so that every *P_i,j_* only depends on the content of 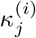. The number of hidden channels are respectively 64, 32 and 16 in our implementation.

### TemplateNet

The TemplateNet *ϕ_t_* encodes a matrix *Q* ∈ ℝ^*b*×*ℓ*^ such that 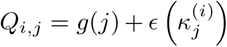, where:

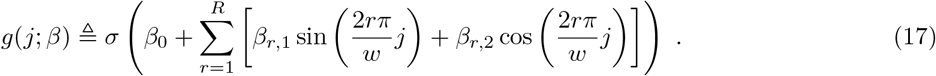

is a sinusoidal function modelled using a truncated Fourier series with amplitude parameters *β*; and *ϵ* is a noise function that outputs positional phase perturbations to the function *g*, parameterized by another convolutional network. We refer to Hoang et al. [8] for explanations of these parameterizations.

### Distance function

The asymmetric distance function proposed by Hoang et al. [8] is given by:

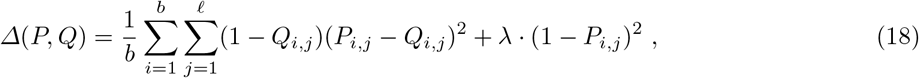

where *P, Q* are the respective outputs of *ϕ_s_* and *ϕ_t_* given *B* and λ is a trade-off constant. Specifically, this distance function corresponds to a weighted ℓ_2_-distance, with larger weights assigned to positions nearer to the template minima, where minimizers are likely to be selected.

## D Details of benchmark sequences

**Table 3.** Descriptions and lengths of sequences used in Section 5.

| Label | Description (Assembly) | Length |
| --- | --- | --- |
| CHRXC | Centromere region of human chromosome X [15] | 3106132 |
| CHR1 | Human chromosome 1 | 233587144 |
| BTR1 | Blautia producta (GCA_004210255.1) | 6354838 |
| BTR2 | Blautia hansenii DSM 20583 (GCF_002222595.2) | 3065949 |
| BTR3 | [Clostridium] scindens (GCA_009684695.1) | 3785527 |
| BTR4 | Blautia producta ATCC 27340 = DSM 2950 (GCA_010669205.1) | 6197116 |

## E Other implementation details

We implement our method using PyTorch and deploy all experiments on a RTX-3080 GPU. Due to limited GPU memory, each training epoch only computes the loss on a randomly sampled batch of 32 substrings of length ℓ = 1500 bases. The conservation component of 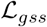 is averaged over 5 random mutations, simulated using a 10% base substitution rate. Evaluation of conservation is likewise obtained using 5 random mutations. Network weights are optimized using the ADAM optimizer [11] with default parameters.

## F Other results

### Effectiveness of training on conservation and density metrics

Fig. 6 demonstrates the individual effect of training the proposed loss 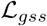 on the conservation and density metrics. We observe that both conservation and density of the syncmer mask are upper-bounded by that of the minimizer mask, which confirms the result of Theorem 1. We observe that 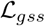 improves conservation but worsens density for the syncmer mask, which is similar to our first experiment. However, this is not the case for the minimizer and complement mask, which obtain significant improvements in both metrics over 600 training epochs. We note that this does not contradict our analysis, as conservation is still bounded by density at any point during the training. Rather, this implies that our method has found a favorable trade-off between the two metrics, which in turn explains the sharper increases in GSS compared to that of syncmer across all experiments.

**Fig. 6.**
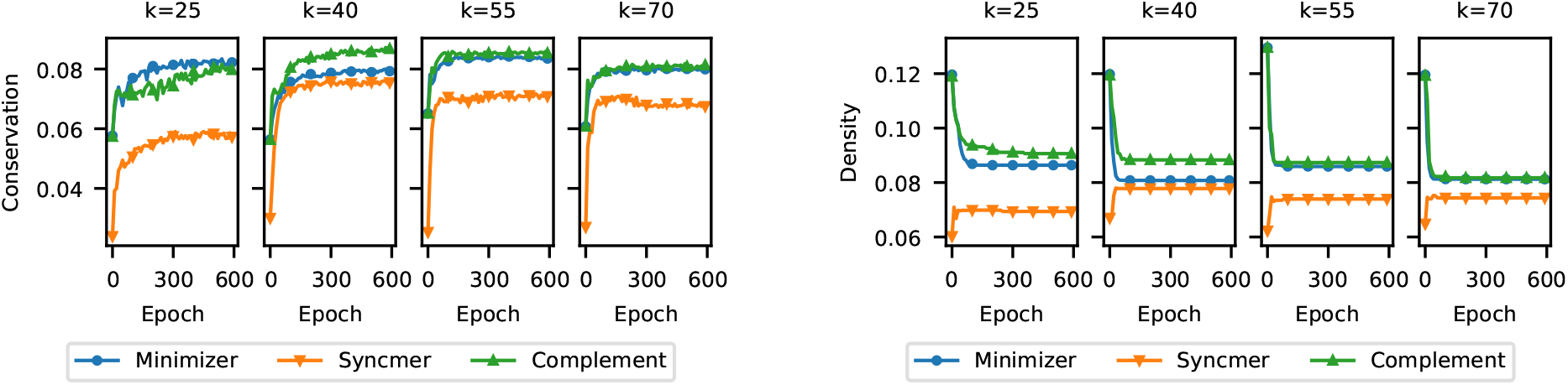
Comparing conservation and density metrics of different masked minimizers vs. number of training epochs on ChrXC with *w* = 15 and *k* ∈ {25,40, 55, 70}.

### GSS profiles of minimizer masks on other bacterial genomes

Fig. 7 shows the scatter plots of all 2^*w*^ masked minimizers trained on Btr1, Btr2 and Btr3 using 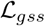 with *w* = 10 and *k* = 15, grouped by the number of 1-entries in their masks. We observe the same increasing pattern of average GSS with respect to number of 1-entries in the mask, thus confirming that the minimizer mask is indeed a good default choice.

**Fig. 7.**
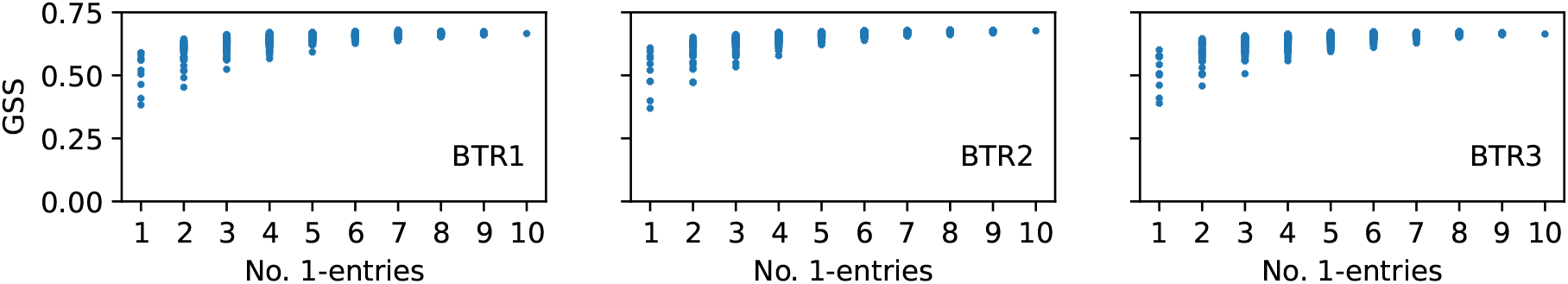
GSS vs. number of 1-entries of all mask minimizers on bacterial genomes Btr1, Btr2 and Btr3.

### Exploiting the relative conservation metric with varying offsets

We repeat this experiment for different syncmer masks with *t* ∈ {6, 7, 8, 9} and plot all results in Fig. 8. In all of these experiments, we observe that the model trained with *n* = 20 sampled mutations per epoch always find the exploit within 1000 – 1500 epochs. The bar plots once again confirm that for each value of *t*, the exploitative solution contains no segment with more than *t* – 1 consecutively decreasing scores. We note that the total count for *t* = 7 is significantly lower than other values of t because the solution contains several segments of monotonically increasing scores that are relatively long, which count towards the > 6 bucket.

**Fig. 8.**
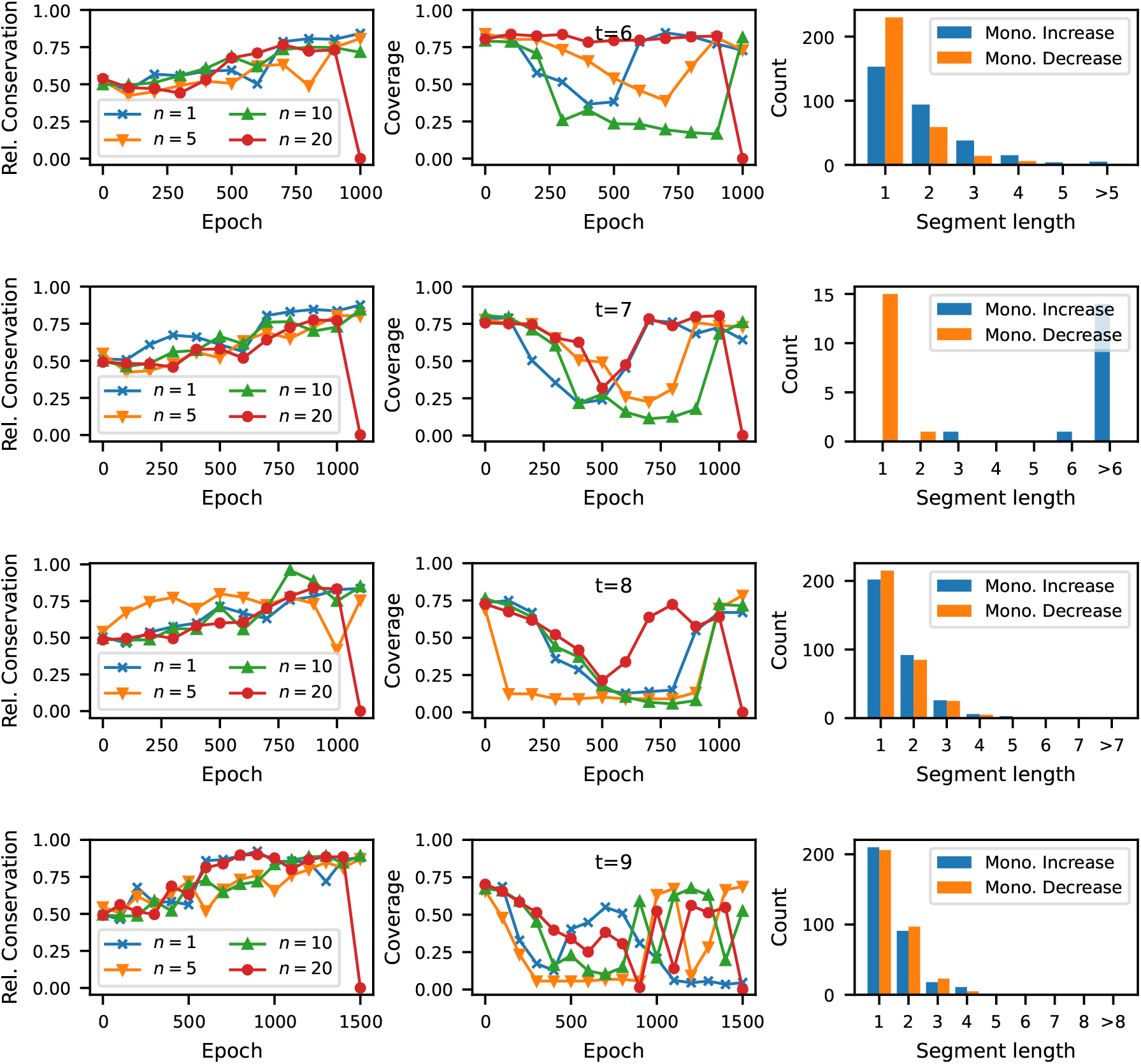
Finding the relative conservation exploit for various syncmer masks with (from top to bottom) offset *t* ∈ {6, 7, 8, 9} with 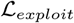, *w* = 10 and *k* = 15.

